# Network Organisation and the Dynamics of Tubules in the Endoplasmic Reticulum

**DOI:** 10.1101/2020.03.11.987891

**Authors:** H. Perkins, P. Ducluzaux, P. Woodman, V. Allan, T. Waigh

**Affiliations:** University of Manchester

## Abstract

The endoplasmic reticulum (ER) is a eukaryotic subcellular organelle composed of tubules and sheet-like areas of membrane connected at junctions. The tubule network is highly dynamic and undergoes rapid and continual rearrangement. There are currently few tools to evaluate network organisation and dynamics. We quantified ER network organisation in Vero and MRC5 cells, and developed a classification system for ER dynamics in live cells. The persistence length, tubule length, junction coordination number and angles of the network were quantified. Hallmarks of imbalances in ER tension, indications of interactions with microtubules and other subcellular organelles, and active reorganisation and dynamics were observed. Live cell ER tubule dynamics were classified using a Gaussian mixture model, defining tubule motion as *active* or *thermal* and conformational phase space analysis allowed this classification to be refined by tubule curvature states.

**STATEMENT OF SIGNIFICANCE:** The endoplasmic reticulum (ER), a subcellular organelle, is an underexplored real-world example of active matter. Many processes essential to cell survival are performed by the ER, the efficacy of which may depend on its organisation and dynamics. Abnormal ER morphology is linked to diseases such as hereditary spastic paraplegias and it is possible that the dynamics are also implicated. Therefore, analysing the ER network in normal cells is important for the understanding of disease-related alterations. In this work, we outline the first thorough quantification methods for determining ER organisation and dynamics, deducing that tubule motion has a binary classification as active or thermal. Active reorganisation and dynamics along with indications of tension imbalances and membrane contact sites were observed.

## INTRODUCTION

The endoplasmic reticulum (ER) is a complex membranous network of tubules and sheet-like regions that is continuous with the nuclear envelope and stretches out to the cell periphery. Narrow tubules (∼100 nm in diameter (1, 2)) intersect at branch points, which typically connect three tubules. The small diameter of the tubules and their relatively rapid dynamics mean that detailed quantitative analysis has only become possible with recent developments in microscopy, such as super-resolution fluorescence microscopy (3). The ER is responsible for the biosynthesis of many proteins and lipids, as well as calcium ion storage and regulation (4–6). Almost all other subcellular organelles interact with the ER at membrane contact sites (MCS) (7), including mitochondria (8–11), endosomes (12– 15), the Golgi apparatus (16) and the plasma membrane (17). Microtubules also interact with the ER via the motor proteins cytoplasmic dynein (18–20) and kinesin-1 (21, 22) and via tip attachment complexes (TACs), which can be formed between the ER and microtubules in the absence of motor proteins (23). EB1, a protein localised to the fast-growing microtubule plus-ends, can bind to STIM1, a protein localised to the ER, to form a TAC (24). This causes a rearrangement of the ER network as the microtubule polymerises. MCS with mitochondria, late endosomes, lysosomes and interactions with microtubules were recently shown to facilitate remodelling of the ER network (25).

The tubules and junctions of the ER network are highly dynamic and oscillate over time (2, 3). The functionality of these oscillations is not yet known, although it has been suggested that mixing of reactants is facilitated by the motion in a *shaken reaction vessel model* (2). This motion is hypothesized to increase the rate of reactions and therefore the rate of protein biosynthesis. Peristalsis of the ER tubules has also been postulated (28), although little quantitative data has yet been presented for this model.

From the perspective of active matter physics (26, 27), the ER represents an important, but relatively unexplored example of active matter. Motor protein activity drives the system out of equilibrium. The tubules are self-assembled from surface active molecules (predominantly lipids, although proteins play an important role in branching and curvature), but resemble polymers at large length scales. The tubules branch and form a complex subcellular network, which is actively driven by molecular motors.

The tubules of the endoplasmic reticulum provide a novel example of driven active polymeric systems that can be compared with more standard systems, such as myosin with actin networks or kinesins with microtubule networks (29–32). A unique feature of the ER is its ability to experience large geometrical and morphological changes due to the elasticity of the lipid tubules e.g. tubules can extend rapidly from the pre-existing network. The ER is therefore a variable non-equilibrium structure that constantly needs to be maintained by active physiological processes.

In this work, the structure and dynamics of the endoplasmic reticulum were studied in detail using fluorescence microscopy of both live and fixed cells. A wide range of image analysis tools were developed including an individual *snakes* analysis of the tubules, Fourier analysis of tubule deformation modes and a *snakes* algorithm to track the junctions. Furthermore, a novel method of specific fluorescent labelling of the ER was developed, based on the genetic constructs designed by Boncompain et al. (33).

In total we quantified the network structure (persistence length, tubule length, junction coordination number and tubule angles), the translational motion of the network, the angular motion of the network, internal fluctuations in the curvature of the tubules (using Fourier analysis) and translational dynamics of the junctions (using MSDs). A classification system for the dynamics of tubules was developed based on these results using a Gaussian mixture model and conformational phase space analysis. Based on this data we refine our previous *shaken reaction vessel model* to be a *kneaded dough model* — large scale geometrical rearrangements constantly happen in the ER due to angular reconstructions combined with active curvature fluctuations on smaller length scales and this may facilitate mixing of protein contents in the ER.

## METHODS

VERO cells, an immortalised cell line derived from the kidney of an African green monkey, were cultured in Dulbecco’s Minimum Essential Media (DMEM) supplemented with 10% Fetal Bovine Serum (FBS, Hyclone, Logan, UT) at 37°C and 8% CO_2_.

### Fixed Imaging

48 hours prior to imaging, Vero cells were cultured on No 1.5 glass coverslips in a 12-well plate. 24 hours prior to imaging, cells were transiently transfected with a construct encoding signal sequence-streptavidin-KDEL (ER-retention sequence), an ER hook from the RUSH system (33). Str-KDEL_neomycin was a gift from Franck Perez (Addgene, Watertown, MA, Addgene plasmid # 65306; http://n2t.net/addgene:65306; RRID:Addgene_65306). This construct allowed subsequent labelling of well-fixed ER using fluorescent biotin, a method which is less susceptible to artefacts than antibody labelling. SS-Str-KDEL DNA (800 ng) was combined with 2 μL of JetPEI (Polyplus Transfection, Illkirch, France) in a total of 100 μL of 150 mM NaCl per coverslip. The solution was left to incubate at room temperature for 15 minutes before being added to the wells.

The next day, the cells were fixed in 3% formaldehyde solution (37 wt.% in H_2_O, 252549, Sigma-Aldrich, St. Louis, MO) with 0.1% glutaraldehyde solution (25% in H_2_O, G5882, Sigma-Aldrich), in PBS for 20-25 minutes at room temperature. Coverslips were washed three times with PBS before a permeabilisation with 0.5% (v/v) Triton X100 in PBS for 10 minutes at room temperature. Unreacted aldehyde groups were then quenched using 1 mg/ml of sodium borohydride in PBS (three five-minute incubations). ATTO 565-biotin (92637, Sigma-Aldrich) was dissolved in DMSO to a concentration of 0.4 mM. After washing in PBS, coverslips were incubated with a 1/2000 ATTO 565-biotin dilution for 25 minutes to label the ER. After three PBS washes were performed, the coverslips were mounted onto glass slides using ProLong Gold (P36930, ThermoFisher, Waltham, MA) and left to cure overnight at room temperature.

Cells were illuminated using a CoolLED pE-300 white excitation system and an appropriate excitation filter. An Olympus BX60, a CoolSNAP EZ camera (Photometrics, Tuscon, AZ) and MetaVue software were used to image the coverslips. Images were imported into Fiji (34). A Gaussian filter with a 1.4 pixel radius combined with a background subtraction with a 2.4 pixel rolling ball radius was used to process the images.

### Live Imaging

The videos of ER dynamics in MRC5 cells and the methods used to obtain them was described previously (2). 48 hours prior to imaging, Vero cells were cultured on glass-bottomed 35 mm dishes (μ-dish, No 1.5coverslip, uncoated, Ibidi, Germany). Cells were transiently transfected with an ER-targeted GFP construct (GFP-LongER), as described in (22), roughly 24 hours before imaging. For each dish, 400 ng of GFP-long-ER and 600 ng of a carrier DNA, pBlueScript, were combined with 4 μL of JetPEI in 100 μL of 150 mM NaCl. This solution was then incubated at room temperature for 15 minutes before being added to the dish.

Before imaging the dishes, pre-warmed phenol red-free imaging media was added to the dishes. This media consisted of Modified Hank’s Balanced Salt solution (with sodium bicarbonate, without phenol red, H8264, Sigma) supplemented with 10% v/v FBS, 2.5% v/v HEPES buffer (1 M, pH 7.0-7.6, H0887, Sigma), 1% MEM non-essential amino acid solution (100x, M7145, Sigma), 2% MEM amino acids (50x, M5550, Sigma), 1% Penicillin Streptomycin (10,000 units/mL, 15140-122, Gibco) and 1% L-Glutamine solution (200 mM, 59202C, Sigma). An Olympus IX71 with a Cairn OptoLED light source and an Olympus PlanApo 100x oil-immersion objective with a numerical aperture of 1.40 were used to view the cells. A standard green fluorescent protein filter set was placed in the emission path. An incubation chamber surrounding the microscope stage was maintained at 37°C throughout the experiment. A Prime 95B sCMOS camera (Photometrics) was used to capture the 400-1000 frame videos with 20-30 ms exposure times and the LED continuously illuminated the sample. Metamorph was used to record the videos.

### Image Analysis and Feature Tracking

The raw videos were imported into Fiji (34). A Fourier transform bandpass filter was applied to the videos in order to highlight structures of a relevant size. Background subtraction using a 2.4 pixel rolling ball radius was then applied. This processing method improved the performance of the tracking algorithm as the signal-to-noise ratio of the images was improved.

In order to track the contours of the tubules over time, a custom modification was applied to the open-source fibre tracking software, FiberApp (35). Given two manually defined points, one at either end of a fibre, FiberApp is designed to find the spatial co-ordinates of a fibre-like object in a single image. An iterative algorithm is used to deform the contour so that it follows the image features. The modification applied to FiberApp requires the user to define the start and end points of the fibre in the first frame, and then uses the tracked start and end points as an input for the next frame. Therefore, the spatial co-ordinates of a tubule can be determined semi-automatically for the duration of a video.

The Fourier modes of each tubule were then calculated using the spatial co-ordinates determined using the modified version of FiberApp. Several custom Matlab scripts based on previous work (36, 37) were written to calculate and analyse the Fourier mode amplitudes.

In order to track junctions in the ER network, a method similar to that described by Xia et al. (38) was used. Videos were initially analysed image-by-image using SOAX, software written to quantify filamentous networks (39). Videos with junctions appearing in greater than 100 frames were processed before tracking began. Figures showing each step in the junction tracking process can be found in Figure S1 in the Supporting Material. Gaps in the initial junction localisation analysis of 10 frames or fewer were accepted. Rosenfeld’s algorithm (40) was used to skeletonise the images and the results were resized. A weighted map of distance was then created of the images. A four-fold increase in the size of the images increased the size of the pattern and therefore the accuracy of the algorithm used to calculate the weighted map of distance. These steps ensured that the maximum intensity occurred at the centre of the tubule while retaining information from the original image.

Junction candidates were then initialised in the neighbourhood of the junction point detected by SOAX, for pixels with intensity greater than zero. A Sobel filter was used to find the intensity gradient and surrounding tubule candidates were detected. Weaker tubule candidates were removed using a non-maximum suppression algorithm and the remaining tubule candidates were linked to the junction. The number of false alarms (*NFA*) was then calculated to find the likelihood of a structure appearing randomly. Junction candidates were only kept if log(*NFA*) < 0. Finally, the junction candidate with the smallest NFA value was taken to be the junction of interest. These two tracking methods (FiberApp and SOAX-based) were used to track the motion of the ER tubules and junctions reliably over whole videos.

## RESULTS

### Network Organisation

The lengths of ER tubules and the angles between tubules at branch points (Figure 1A) were measured in fixed Vero and live MRC5 cells. Tubules in the ER networks were found to be of the order of micrometres in length, with a mean length of 1.072 ± 0.009 *μ*m for live MRC5 cells (n = 4521 tubules, mean ± SEM) and 1.120 ± 0.007 *μ*m for fixed Vero cells (n = 6537 tubules, mean ± SEM). A histogram of tubule lengths is shown in Figure 1B. When the contour lengths of tubules were studied in live cell videos for both MRC5 and Vero cells, lengths were found to be fairly constant over short time scales, with the exception of a small number of rapid extension events (an example of which can be seen in Figure 1B, inset).

**Figure 1.**
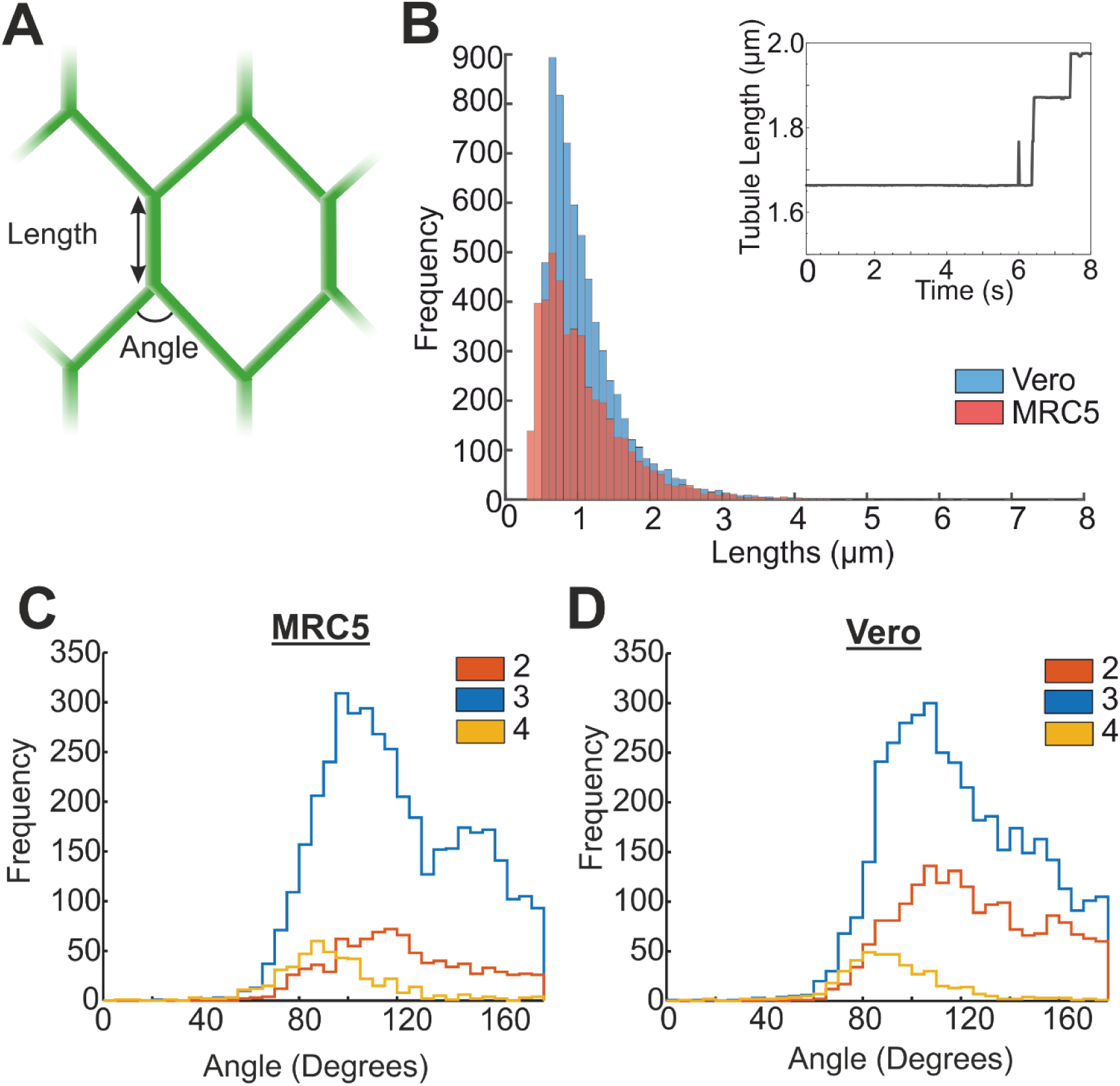
Lengths of ER tubules and angles subtended between tubules at junction points. **A** Schematic diagram of a tubule within the network showing the lengths and angles measured. **B** Histogram of Vero (blue) and MRC5 (red) ER tubule lengths, measured in fixed Vero cells and stills of live MRC5 cells. The mean length was 1.120 ± 0.007 *μm* for Vero cells (n = 6537 tubules, mean ± SEM) and 1.072 ± 0.009 *μm* for MRC5 cells (n = 4521 tubules, mean ± SEM). Inset: Example of rapid extension of a tubule. At small timescales, this tubule also shows small fluctuations in length. **C** Angles measured in the ER network of MRC5 cells, separated by the co-ordination number of the junction. **D** Angles within the ER network of Vero cells, separated by junction co-ordination number.

The average angles subtended by tubules at branch points were found to vary with the number of tubules connected in both cell types (Figure 1C and D). Combining the results from both cell types, the ER network was found to have junctions consisting of either 2, 3 or 4 tubules, with 3-way junctions being the largest population (69 ± 2 %) (41), followed by 2 (24 ± 2 %) and then 4 (8 ± 2 %). For junctions between four tubules, the mean angle measured between tubules was 93.2 ± 0.8°. This corresponds to an equilibrium configuration of the branch point. It is worth noting that with this method of analysis, it is impossible to discern whether the membrane of these tubules has fused to form a branch point or if the tubules are simply overlapping. A bimodal angular distribution for 3-way branch points (more pronounced in MRC5 cells, blue in Figure 1C) was observed with 103° ± 1° observed roughly twice as frequently as 153° ± 3°. Under symmetric equilibrium conditions, a separation of 120° is expected, with equal tension in all 3 tubules. However, branch points may each have two 100° angles and one 160° angle if there are heterogeneous non-equilibrium forces, and therefore tensions, within the network. As the ER is highly dynamic and undergoes constant rearrangements, it seems likely that the majority of tubules experience such non-equilibrium conditions. The angles observed between tubules at 2-tubule branch points ranged from ≈ 20° to less than 180°. Given that a branch point between two non-colinear flexible fibres cannot be formed without a third component to balance the forces, this result indicates that the ER forms membrane contact sites with other cellular structures or interacts with microtubules to create these branch points.

### Tubule Persistence Length

Fluorescence images of live cells were used to calculate the persistence length of ER tubules in Vero cells, using a similar method to Georgiades et al. (2). The persistence length gives a measure of the flexibility of fibre-like objects. It is defined as the distance along the contour of the fibre over which angular correlations in the tangential direction decorrelate. Fibres with lengths much greater than their persistence lengths are classified as flexible, whereas the opposite is true for rigid fibres. Semi-flexible fibres are those with comparable lengths and persistence lengths.

FiberApp (35) was used to trace the contours of ER tubules in images of live Vero cells (see Methods). Tubules were defined as the sections of the network between two branch points. The contour lengths, *L*_*c*_, and end-to-end distances, *R*, were determined. The mean square end-to-end distance of a fibre, ⟨*R*^2^⟩, can be related to the contour and persistence length, *L*_*p*_, as (42)

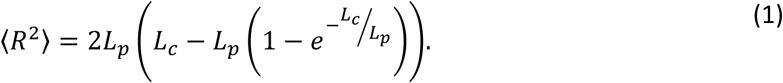

Fitting the values of *L*_*c*_ and *R* to Eq. 1 led to a persistence length of 8.2 ± 0.2 *μ*m for Vero cell ER tubules (mean ± SD). This value is in reasonable agreement with the persistence length of 4.71 ± 0.14 *μ*m previously found for fixed and live MRC5 cells (2) and again indicates that ER tubules can be modelled as semi-flexible filaments. The larger persistence length measured for Vero ER tubules also indicates that ER tubules may be under greater tension in Vero cells than in MRC5 cells.

### Tubule Dynamics

ER tubule dynamics were studied using mean squared displacements (MSDs), a standard method of quantifying particle dynamics. The MSD is defined by

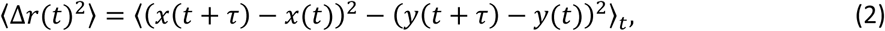

where *x*(*t*) and *y*(*t*) are the co-ordinates at time *t* and *τ* is the lag time. MSDs of lateral tubule movements were calculated by tracking the position of a point on a tubule over time. A line was drawn perpendicularly to the tubule and the co-ordinates of the tubule on this line were tracked, similarly to (2), but with the inclusion of a background subtraction step. As longitudinal tubule motion was not measured in this study, and generally has a much smaller amplitude than transverse motion, the MSD can be assumed to describe the transverse motion of the tubule, ⟨Δ*r*(*t*)^2^⟩ = ⟨Δ*r*_⊥_(*t*)^2^⟩ + ⟨Δ*r*_‖_(*t*)^2^⟩ ≈ ⟨Δ*r*_⊥_(*t*)^2^⟩ (26). The transverse MSD can be approximated as a power law: ⟨Δ*r*_⊥_(*t*)^2^⟩ ∝ *τ*^*α*^, in which the exponent, *α*, describes the motion of the tracked object. Brownian motion is described by *α* ≈ 1, whereas super-diffusive motion has an exponent 1 < *α* < 2, and 0 < *α* < 1 describes sub-diffusive motion. For semi-flexible fibres oscillating in a viscous medium with no applied tension, ⟨Δ*r*_⊥_(*t*)⟩ ∝ *τ*^3/4^ is expected, whereas for tension-dominated regimes theory predicts that ⟨Δ*r*_⊥_(*t*)⟩ ∝ *τ*^1/2^ (26).

Prior to fitting a power law to the MSDs, a constant offset was removed from each measured MSD, ⟨Δ*r*(*t*)⟩_*m*_. This increased the time scale over which the power law could be fitted to the MSDs and minimised the effect of static errors due to experimental signal-to-noise limitations (43). A straight line fit was applied to the first 5 points of the MSD and the resulting intercept, *σ*, was subtracted from the measured MSD (44, 45),

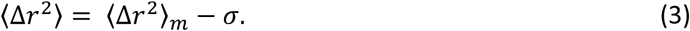

The raw MSD data is shown in Figure 2A, and Figure 2C displays the same data with the constant offset removed using Eq. 3. The distribution of *α* values for the MSDs in Figures 2A and 2C are displayed in Figure 2B and 2D respectively. Many of the raw MSDs were found to have an exponent of the order of 0. However, very few of the corrected MSDs were had *α* < 0.2. This indicates that the section of the MSDs dominated by static error is reducing the exponent determined. The corrected exponents show that the majority of tubules moved sub-diffusively, with *α* < 1. However, a significant proportion (29 ± 1%) of the tubules displayed super-diffusive motion associated with active, motor-driven motility. These results are in agreement with our previous work (2).

**Figure 2.**
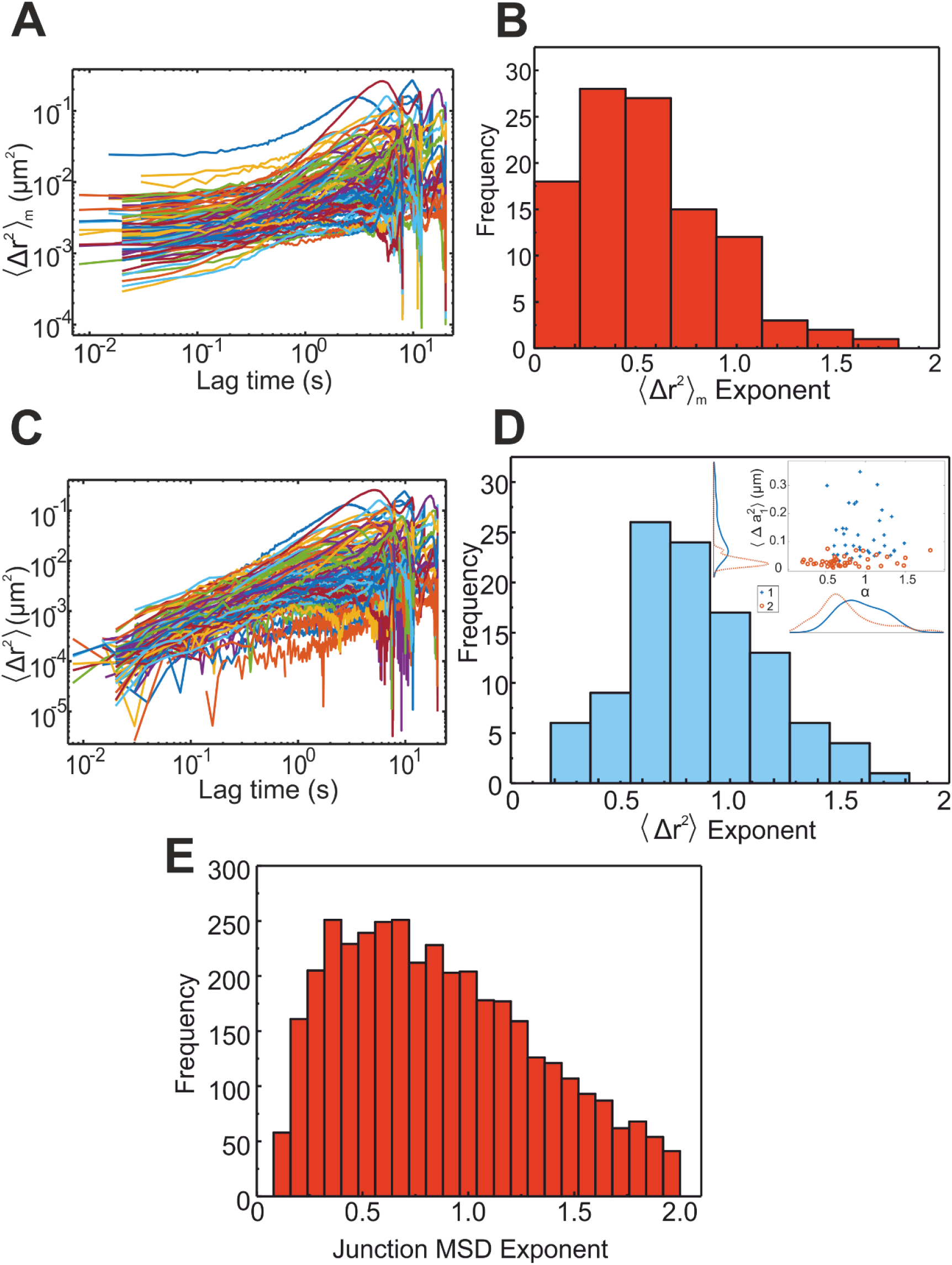
Mean squared displacement of transverse motion of the ER in live Vero cells. **A** Measured transverse MSDs of tubules. Traces begin and end at different time points due to the slight variation in frame rates and total times of the videos recorded. **B** MSDs were fitted with a power law, τ^α^. The exponents of the MSDs in **A** are depicted. **C** The initial constant section of the measured MSDs shown in **A** was subtracted to extend the fitting region and therefore give a more accurate estimation of α. The resulting MSDs are depicted. **D** The power law exponents of the corrected MSDs shown in **C**. Inset: Marginal probability distributions of the Gaussian mixture model used to cluster tubule dynamics. A peak in the marginal of one distribution is observed at *α* ≈ 0.5. This result is in agreement with Georgiades et al. (2) and the theoretical prediction ⟨Δ*r*_⊥_(*t*)⟩ ∝ *τ*^1/2^ (26). **E** MSD power law exponents for 3763 tracked junctions of the ER network in live Vero cells. The mean exponent found was 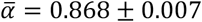 (mean ± SEM).

### Junction Dynamics

The junctions of the ER, as well as the tubules, are known to oscillate (46). Similarly to tubules, the junctions of the ER network can be created or destroyed due to the rearrangement dynamics of the network. New junctions can form when a tubule extends and fuses with another region of the network and ring closures can merge multiple junctions. Forces within the network can also cause tubules at junctions to dissociate, destroying a junction, although this is rare. In this work, branch points were tracked using videos of the ER network in live Vero cells.

The coordinates of the junctions over time were used to calculate MSDs. As for the ER tubules, Eq. 3 was used to remove the constant offset. Figure 2E shows the power law exponents for 3763 tracked junctions in live Vero cells. Both super-diffusive and sub-diffusive junctions were observed in the network, with the mean exponent being sub-diffusive 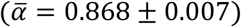. Therefore, the majority of junctions between tubules in the ER network oscillate sub-diffusively, with a significant population of super-diffusive junction motions which are actively driven. ER junction dynamics in HeLa cells were previously tracked by Speckner et al., who concluded that junctions moved with *α* ≈ 0.5 (46), in agreement with our conclusion that junctions oscillate sub-diffusively.

### Mean Squared Angles

The angular dynamics of tubules and the local geometry changes of the network were also studied by calculating the mean squared difference of the angle between the ends of the ER tubules, MSΘ. Specifically, Θ, the end-to-end angle for a tubule in each frame was determined (Figure 3A) and the mean squared difference in Θ was then found using

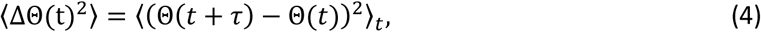

where *t* and *τ* are time and lag time respectively. As for tubule and junction MSDs, the constant offset was removed from the data and a power law was fitted, with 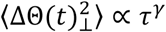. Previous work by LeGoff et al. (47) showed that angular fluctuations in single F-Actin filaments *in vitro* followed a *τ*^1/4^ power law. Contrary to this observation, almost all tracked angular fluctuations of live Vero ER tubules had a larger exponent (Figure 3B, C). A *τ*^0.63±0.02^ power law was observed, with a small population of tubules displaying super-diffusive angular dynamics. Given that the persistence length of actin is ∼17 *μ*m (47–50), whereas ER tubules have *L*_*p*_∼8 *μ*m (2), it may be expected that ER tubules exhibit more dynamic angular fluctuations due to their increased flexibility. However, it is surprising that tubules attached in a network have larger angular fluctuations than fibres that are free at both ends. This result again indicates that the ER network is dynamic at tubule ends where junctions occur, in addition to transverse tubule fluctuations.

**Figure 3.**
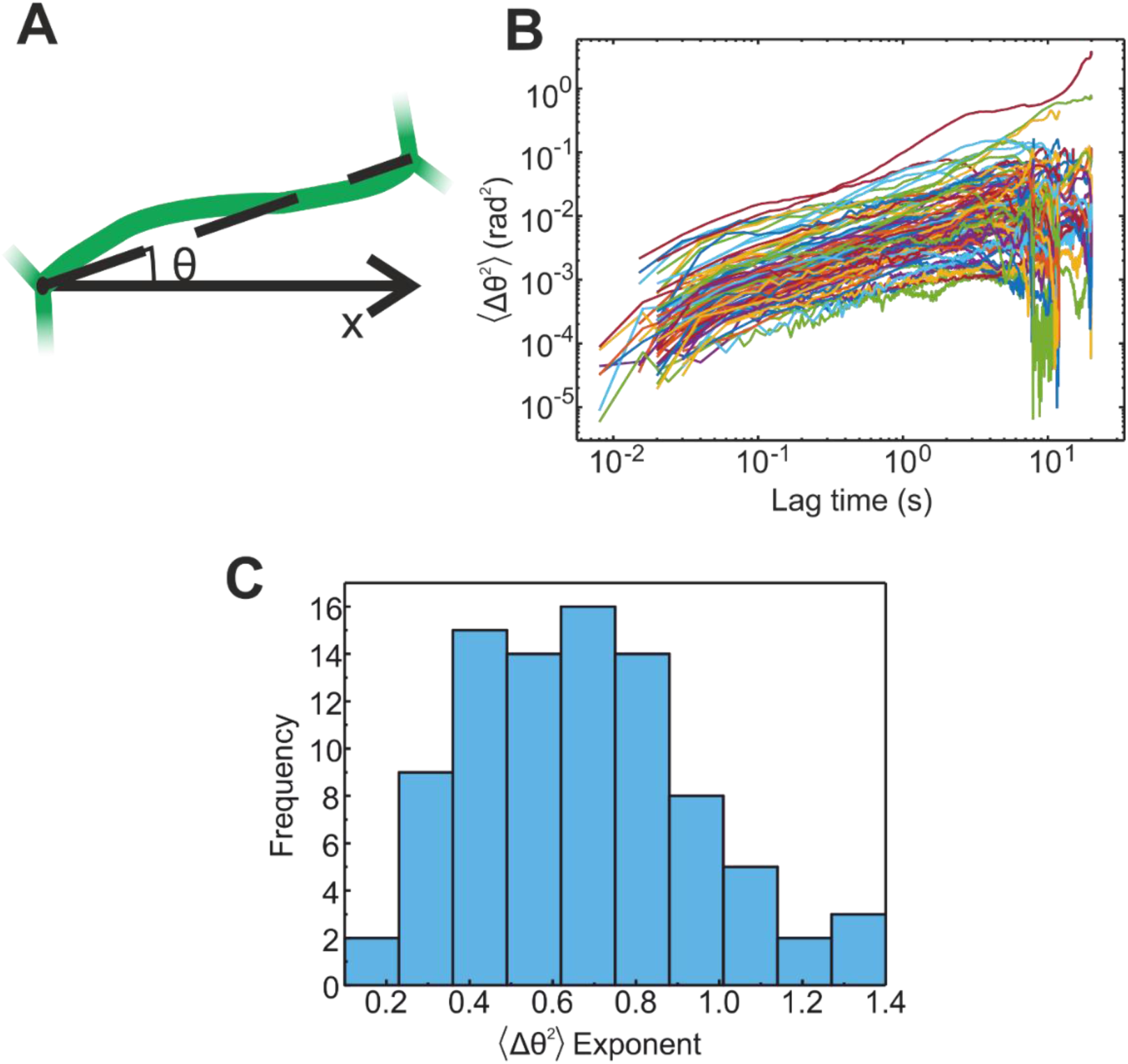
Mean squared angles subtended between the ends of ER tubules (MSΘ) in live Vero cells. **A** Schematic diagram showing the end-to-end angle, Θ, of a tubule (green) with respect to the arbitrarily chosen x-axis. **B** MSΘs for live Vero cells as a function of lag time. As in Figure 2C, the initial constant region has been removed. **C** Histogram of the power law exponents for the corrected MSΘs.

### Analysing Tubule Curvature using Fourier Mode Decomposition

In order to study the curvature dynamics of ER tubules, tubule contours were decomposed into Fourier modes (37, 49). This led to a more detailed description of individual tubule dynamics. Contours were traced for each frame of a video, using a version of FiberApp (35) modified for this work (see Methods). Two examples of tracked tubule contours are shown in Figure 4A. The co-ordinates of the tubule at the *k*^th^ point along the backbone of the tubule, (*x*_*k*_, *y*_*k*_), were used to find the tangent angle, *θ*_*k*_, at each point,

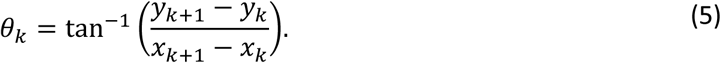

**Figure 4.**
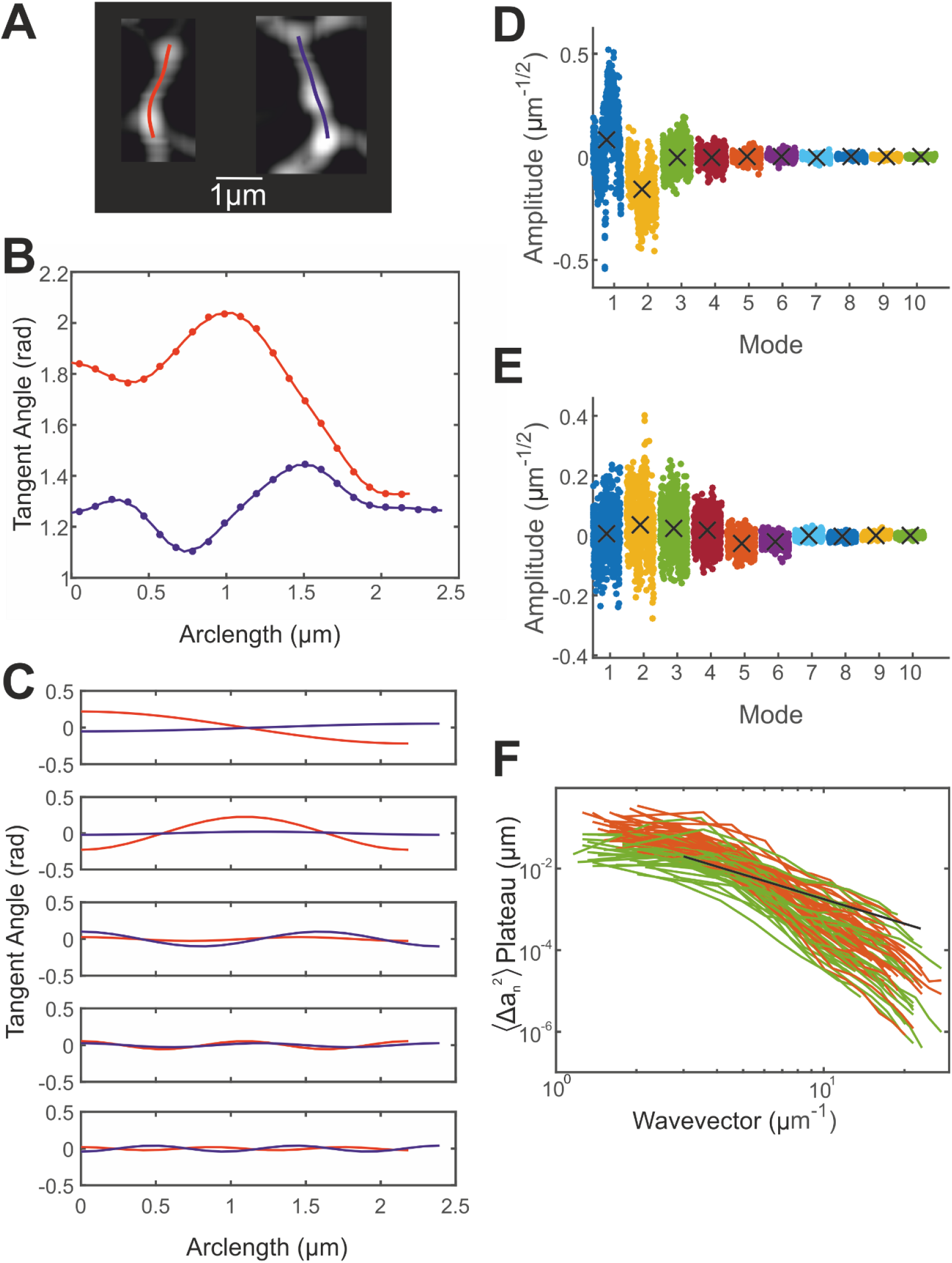
Tubule contours are modelled with Fourier modes. **A** Fluorescence microscopy images of tubules in live Vero cells with tracked contours overlaid. **B** Tangent angles as a function of arclength, *θ*(*s*), for the tubules in **A**, as calculated from the tracked points along the tubules using Eq. 5, are depicted using a solid line. The colours correspond to the tubules shown in **A.** The tangent angles calculated using Eq. 8 with 10 Fourier modes to model the contour are depicted using points. **C** The first 5 Fourier modes used to describe the tubules are shown, with the first mode displayed in the upper panel and the mode numbers increasing to 5 for the lowest panel. The larger mode amplitudes of the red tubule are due to its more curved contour. **D** Fourier mode amplitudes for the tubule depicted in red in **A**, shown for all frames of the video. Black crosses show the mean mode amplitudes. This tubule has non-zero average curvature, seen in the non-zero mean mode amplitudes of the first two modes. **E** Fourier mode amplitudes for the tubule depicted in blue in **A**. In this case, the means are approximately zero, indicating zero average curvature. **F** Plateau values of the mean squared difference in Fourier mode amplitudes plotted against the wavevector. The ensemble average fit to Eq. 10 is shown in black. The deviation from this fit at larger wavevectors is expected for semi-flexible fibres deformed by active influences (such as motor proteins) (29), as opposed to thermally oscillating fibres.

Each tubule was segmented into smaller lengths along its contour as

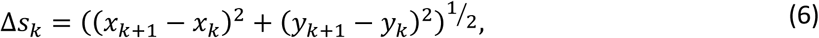

with the arclength at the *k*^th^ point along the tubule, *s*_*k*_, then defined as a summation of the segments:

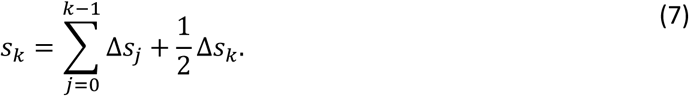

The shape of each tubule was also expressed as a sum of Fourier modes. As the tubule ends are not free, but rather fused to other tubules in the network, cosine as opposed to sine modes were chosen. An infinite sum of modes, *θ*_*n*_(*s*), each with a mode number, *n*, and amplitude *a*_*n*_, fully described the shape at each point of the fibre, *θ*_*s*_,

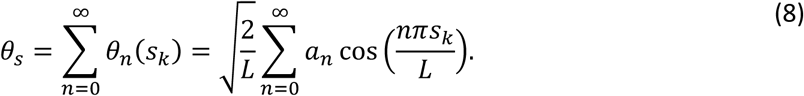

In Eq. 8, *L* is the length of the tubule and each mode is normalised by 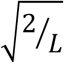, such that modes are length independent. Mode amplitudes were found using the approximation that for a sufficiently large number of modes (*n* = 10 was used in this work),

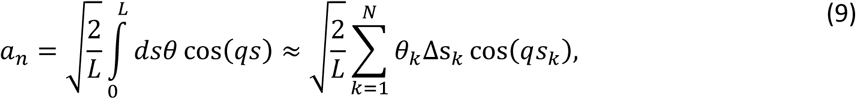

where *N* is the number of segments that the tubule was split into and the wavevector, *q*, is equal to *nπ*/*L*. A script was written in Matlab to decompose the tracked contours of the tubules into Fourier modes by finding the tangent angle, *θ*_*k*_, at each tracked point along the arclength of the tubule, *s*_*k*_. Figure 4B shows *θ*_*k*_ as a solid line and *θ*_*s*_ as points, calculated using 10 Fourier modes, along the arclength of the example tubules. The first 5 modes used to model each of the contours are shown in Figure 4C.

The average curvature in ER tubules was also found using Fourier modes amplitudes. For a rigid, straight fibre *a*_*n*_ would be roughly 0 over time for all *n*, as no fluctuations would occur. However, a curved fibre oscillating around its curved equilibrium state would have non-zero *a*_*n*_, with mean mode amplitudes indicating the equilibrium curvature of the fibre. Tubules were classified into 2 groups using the Fourier mode amplitudes. Tubules for which greater than 1.5*σ* (>93% of the points) of *a*_*n*_ values fell on one side of zero for at least one Fourier mode, *n*, were said to have sustained curvature. If this condition was not met, no sustained curvature was found. Figure 4D shows the variation of each Fourier mode amplitude with time for the tubule shown in red in Figure 4A, which oscillates with persistent curvature. The blue tubule in Figure 4A has no sustained curvature, as shown by the mean mode amplitudes of zero in Figure 4E. Contact sites between the ER and other cellular structures may facilitate these curved tubule configurations, also known as states of prestress (36, 37).

The Fourier mode amplitudes for pairs of modes also provide information about tubule curvature over time. Transitions between states, 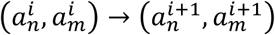, where 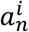 denotes the amplitude of mode *n* in frame *i*, were interpolated and plotted in 2D coarse-grained conformational phase space (31, 51, 52). Figure 5A shows example phase space plots of the first two modes for various tubule conformations: *straight* (Figure 5Ai), *sustained curvature* (Figure 5Aii) and *transient curvature* (Figure 5Aiii and iv). Note that the axes scales vary between subpanels.

**Figure 5.**
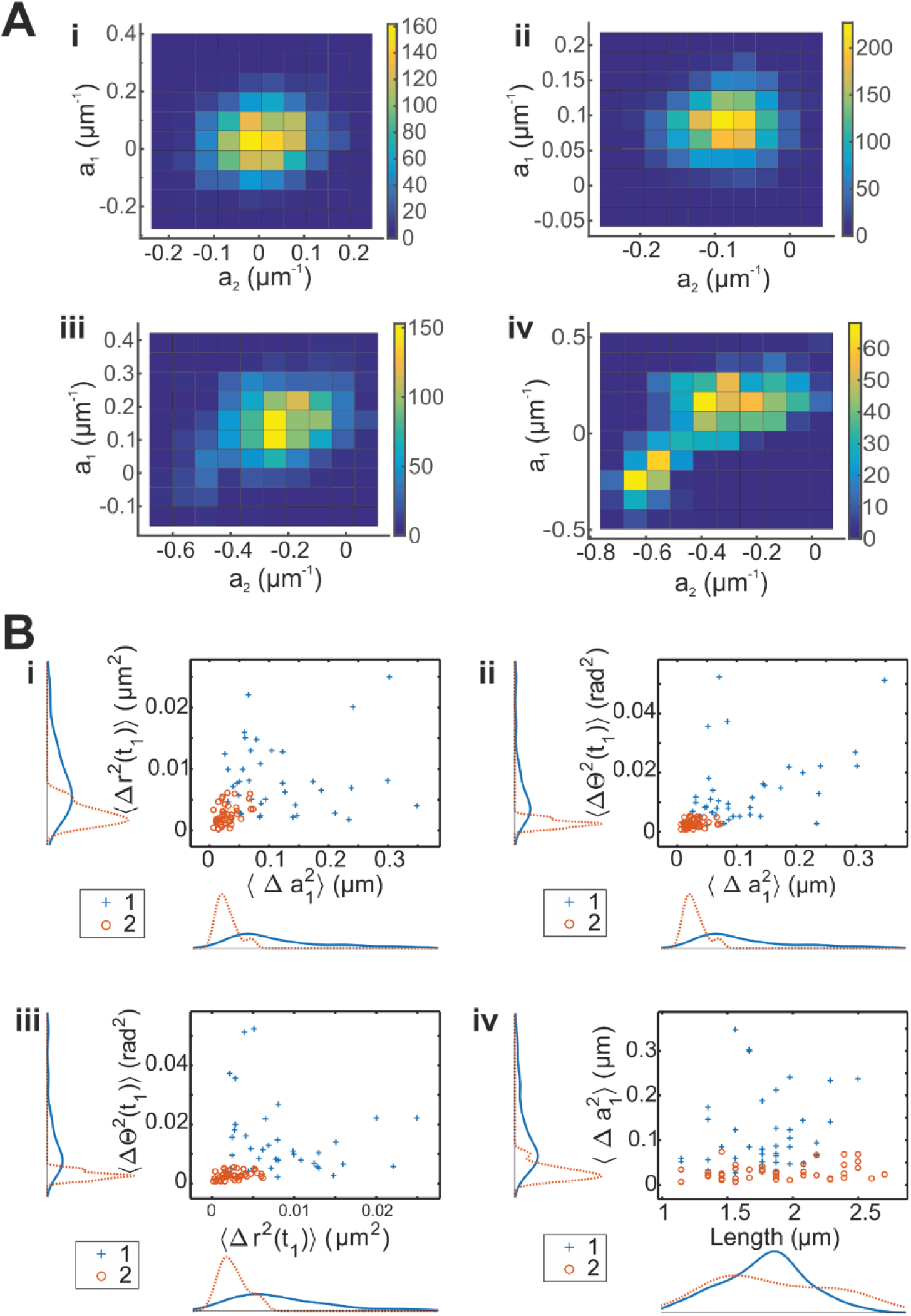
Coarse-grained configurational phase space plots and Gaussian mixture models calculated for the tubules’ dynamics. **A** Configurational phase space plots for the amplitudes of the first and second Fourier modes, *a*_*1*_ and *a*_*2*_. Four examples are shown – (i) modes centred at zero, (ii) modes centred at a non-zero point, (iii) comet-shaped modes and (iv) modes with two separated centres. Note that axis scales are different in each subpanel. Tubules with negligible curvature are expected to have shapes such as (i), whereas (ii) corresponds to tubules with constant, sustained curvature. Configurations (iii) and (iv) represent transient changes in curvature, with very short-lived changes depicted by small changes from circular shapes, as in (iii). Longer-lived deformations lead to more frequent deviations from circular shapes, as in (iv). **B** Marginal probability distributions of the Gaussian mixture model used to classify tubule motion as *active* or *thermal*. Nominal MSD values, nominal MSΘ values, first Fourier mode amplitudes and lengths are plotted. Tubules in the active population, population 1, are shown with blue crosses in all four subpanels, whereas thermal tubules are shown with orange circles. Active tubules tend to have larger MSD, MSΘ and 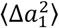 plateau values, as seen in (i-iii). The length-independence of the populations is shown in (iv).

The presence of circulatory currents in conformational phase space due to the breakdown of detailed balance is another signature of active, non-equilibrium dynamics (31, 51, 52). However, due to the fast dynamics of ER tubules, precise currents in phase space could not be determined and the circulations calculated were not statistically significant (verified with Monte Carlo simulations, data not shown).

### Dynamic Persistence Length

The mean squared difference in Fourier mode amplitudes, ⟨Δ*a*_*n*_^2^⟩, provided another estimate of the persistence length of semi-flexible fibres (see Fig. S2 in the Supporting Material). Theory predicts that for thermal oscillations, the plateaus in the MSDs of Fourier modes, ⟨Δ*a*_*n*_^2^⟩_*plat*_, should be inversely proportional to the square of the wavevector, *q* (49, 50),

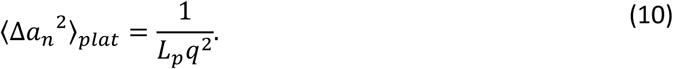

However, recent theoretical work by Weber et al. (29) predicts that for a semi-flexible fibre deformed by active cross-linkers, Eq. (10) will no longer hold for large wavevectors. Instead, a deviation from this pattern will occur, with ⟨Δ*a*_*n*_^2^⟩_*plat*_ becoming smaller than expected for large *q* values. However, Weber et al. found that the data did follow *q*^−2^ for a sufficiently large range in *q* to find an estimate for *L*_*p*_. Using Eq. (10), the persistence length found was 5.6 ± 0.7 *μ*m (n = 83 tubules, mean ± SEM, see Figure 4F). Wavevector values of less than 3 *μ*m^−1^ were not included in the fit, as these modes are suppressed due to the finite separation of cross-linkers (MCS and interactions with microtubules in this model) along the fibre (29, 37). The deviation from the expected *q*^−2^ dependence indicates the presence of active motors and organelles (29). The *L*_*p*_ calculated is reasonably close to the geometrical estimate found using Eq. (1), 8.2 ± 0.2 *μ*m. The difference in values is probably due to the unaccounted effects of dynamics in the geometrical measure and we would expect the Fourier mode method to be more accurate because the range of *q* can be constrained to reduce the effects of activity.

### Gaussian Mixture Model Clustering

To investigate whether patterns were present in the measured tubule dynamics, machine learning techniques were used as they provide useful, automatic, non-biased tools to classify phases in a diverse range of data sets (53). A Gaussian mixture model was used with the following variables: tubule length, L, MSD exponent, α_MSD_, MSΘ exponent, γ_MSΘ_, the values of the MSD and MSΘ at *t*_1_ = 1s, ⟨Δ*r*^2^(*t*_1_)⟩ and ⟨ΔΘ^2^(*t*_1_)⟩, the first Fourier mode amplitude plateau ⟨Δ*a*_1_^2^⟩_*plat*_ and a measure of tubule bending activity, β. To find β, Eq. (10) was modified such that ⟨Δ*a*_*n*_^2^⟩_*plat*_ ∝ *q*^−*β*^ and the data in Figure 4F was fitted.

Figure 5B shows the length-independent separation of the tubules into two clusters. Very little separation between the two populations was observed for the variables not shown in Figure 5B (see Fig. S3 in the Supporting Material). Tubules in the first population are dynamic, with large fluctuations in *a*_1_ and greater ⟨Δ*r*^2^(*t*_1_)⟩ and ⟨ΔΘ^2^(*t*_1_)⟩ values than the second population. Together, these results indicate a separation of tubules into those with dynamics more similar to active motion (population 1) and those that lack large scale rearrangements and seem to be oscillating passively (population 2). ER tubules can therefore be classified into two populations of dynamic motion: *active* or *thermal*.

## DISCUSSION

From a combination of static and dynamic experiments we can begin to form a quantitative model for the endoplasmic reticulum. On the level of single tubules they act as semi-flexible polymers with mean contour length 1.120 ± 0.007 *μ*m and persistence length 5.6 ± 0.6 *μ*m, determined using Fourier analysis, in Vero cells. The majority of junctions connect 3 tubules, although their angular distributions indicate that tubules experience different applied forces. Mechanical stresses in the tubules have multiple signatures in our experiments, including asymmetrical angles at junctions, MSDs with exponents less than ¾ (54), sudden extensions of tubules and non-zero curvatures (states of prestress).

Structural links between the ER and other subcellular organelles or cytoskeletal structures such as microtubules may account for junctions in the ER consisting of only 2 tubules. CLIMP-63, a transmembrane protein, has been shown to form static links between the ER and microtubules (55, 56). CLIMP-63 is resident in the ER and has a cytoplasmic tail that facilitates binding of the ER directly to microtubules. REEP-1, a protein that localises to tubular ER has also been shown to mediate microtubule-ER interactions (57). Overexpression of REEP-1 caused ER tubules to align with microtubules, indicating that it acts as a linking protein between the two organelles. These connections may also play a role in the positioning of the ER within the cell.

Careful analysis of mean squared displacements of tubules and junctions indicates that the motion of the ER network is predominantly sub-diffusive. Therefore, the majority of the ER network is oscillating passively due to the tension under which it is held and the motion of the surrounding cytoplasm and organelles. Junction MSD exponents are in reasonable agreement with previous results for HeLa cells (46), although a slightly more sophisticated algorithm for junction tracking was implemented in our work. Super-diffusive dynamics were observed for a small population of both tubules and junctions. This directed motion may be caused by motor proteins (22) or by contact sites between the ER and other motile organelles, such as endosomes or mitochondria (11, 12).

Fourier mode decomposition of the tubules revealed states of sustained curvature, or prestress, as well as transient curvature, possibly caused by membrane contact sites with other organelles or interactions with the cytoskeleton. A systematic deviation from the *q*^−2^ distribution for the Fourier mode plateaus expected for thermalized dynamics indicates that active linkers, such as motor proteins, play a role in ER dynamics (29).

The tubules have surprisingly large angular fluctuations 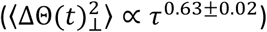 when compared with the expected angular dynamics of non-cross-linked semi-flexible polymers, ⟨*θ*^2^⟩∼*τ*^1/4^ (26), and those observed in free F-Actin (47). Cross-linkers and the network configuration of tubules may contribute to the highly variable geometry observed in the ER network.

A classification system for tubule dynamics was developed using a Gaussian mixture model and Fourier mode decomposition. Tubules were classified as *active* or *thermal* and sub-classified as *curved, transiently curved* or *straight* using this framework. These classifications can be related to the ER network in a live cell, shown in Figure 6A, where a *straight thermal* tubule and an *active transiently curved* tubule are highlighted in (i) and (ii) respectively. Figure 6B displays a diagram of all possible dynamic states of ER tubules that can be detected using this sub-classification scheme.

**Figure 6.**
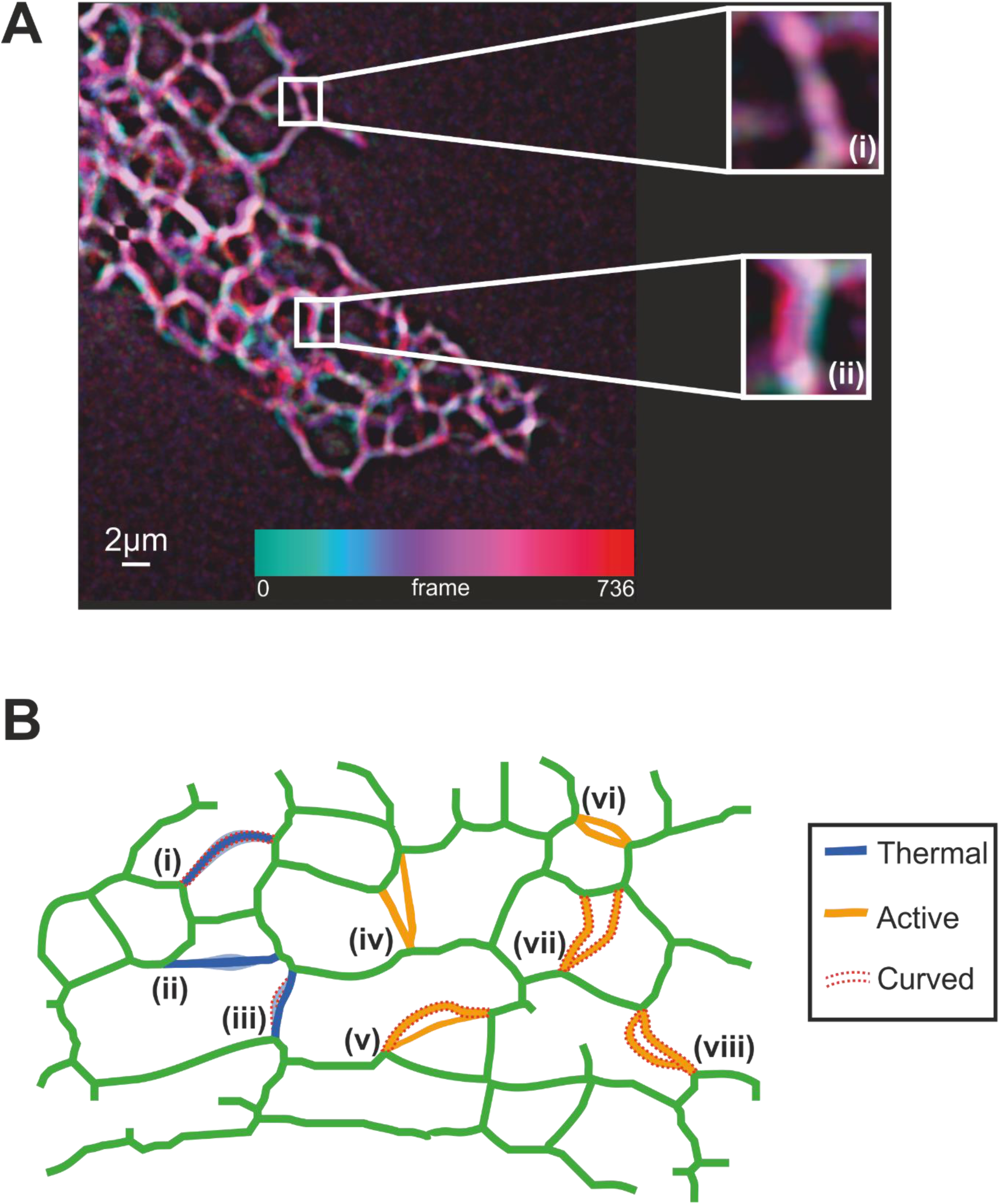
ER tubule classification. **A** Time series overlay of a network section in a live Vero cell. Colours correspond to the frames in which the ER is detected in a certain conformation. Some tubules are relatively stationary throughout the video (as seen in (i), a *thermal straight* tubule) whereas other tubules are more dynamic and undergo curvature changes (as seen in (ii), an *active transient* tubule). **B** Schematic diagram of the sub-classification of ER tubules as *straight, curved* or *transiently curved*. Tubules classified as *active* (shown in orange, (iv)-(viii)) have more significant fluctuations or rearrangements, whereas tubules classified as *thermal* (shown in blue, (i)-(iii)) exhibit smaller amplitude movements. A red dashed outline indicates tubule curvature. *Active* tubules that are *straight* or *curved* can undergo large-amplitude fluctuations (as in (vi) and (viii) respectively), the junctions can undergo large rearrangements (as in (iv) and (vii) respectively) or high degrees of curvature may be observed in some frames (as in (v)). *Thermal* tubules may have be *curved*, as depicted in (i), *straight*, as in (ii) or *transiently curved*, as in (iii). Tubules with transient curvature are curved for a portion of the tracking, but not the entirety.

Along with the new methods developed to describe the organisation of the ER network, this classification system facilitates the quantification of the ER and allows us to refine our previous reaction vessel model into a *kneaded dough* (KD) model. The KD model postulates that large scale coherent motion of the tubules increases the reaction kinetics and is a combination of tubule shaking (changes in curvature and translational motion), geometrical organisation (angular fluctuations) and rapid tubule reorganisation (sudden growth and retraction of tubules). In the future it is hoped this behaviour can be explored in a wider range of cell types and that a rigorous link between ER dynamics and the kinetics of protein production can be made.

## CONCLUSION

We developed a new method to fluorescently label the ER in eukaryotic cells. Image analysis of fluorescence microscopy data allowed us to develop a detailed picture of the ER tubule network and dynamics. A classification system was developed for ER tubule dynamics, based on a Gaussian mixture model using the Fourier mode amplitudes, MSDs and angular fluctuations, which allowed the tubules to be classified as either *thermal* or *active*. Based on the occurrence of prestresses in the tubule curvature, the active and thermal classifications could be further categorised as *straight, curved* or *transient*. Together, these analytical tools provide a new detailed, quantitative description of the endoplasmic reticulum in live cells whose activity can be described by a kneaded dough model i.e. both tubule shaking and tubule rearrangement are important to increase reaction kinetics.

## AUTHOR CONTRIBUTIONS

H.P. performed the experiments, developed the tubule tracking algorithms and analysed the data. H.P., T.W. and V.A. wrote the manuscript. P.D. developed the junction tracking algorithm and analysed the data. T.W., P.W. and V.A. designed the research.

## ACKNOWLEDGEMENTS

We would like to thank the EPSRC for funding this work under the PhD studentship of H.P. and the MRC for funding the construction of the STORM microscope (grant code EP/F062966/1). We gratefully acknowledge the Henry Royce Institute for funding the purchase of the adaptive optics device. Furthermore, we would like to thank Drs Pantelis Georgiades and Henry Cox for their help in developing analysis methods, Dr Mark Johnston for his assistance with live cell imaging and Johanna Blee for useful discussions.

